# Efficient RNA-mediated reprogramming of human somatic cells to naïve pluripotency facilitated by tankyrase inhibition

**DOI:** 10.1101/636670

**Authors:** Nicholas Bredenkamp, Jian Yang, James Clarke, Giuliano Giuseppe Stirparo, Ferdinand von Meyenn, Duncan Baker, Rosalind Drummond, Dongwei li, Chuman Wu, Maria Rostovskaya, Austin Smith, Ge Guo

**Affiliations:** Wellcome–MRC Cambridge Stem Cell Institute, University of Cambridge, Cambridge CB2 1QR, United Kingdom; Department of Biochemistry, University of Cambridge, Cambridge, CB2 1GA, United Kingdom; Guangzhou Institutes of Biomedicine and Health (GIBH), Chinese Academy of Sciences, Guangzhou, China, 510530; Department of Medical & Molecular Genetics, King’s College London, London SE1 9RT, United Kingdom; Institute of Food, Nutrition and Health, ETH Zurich, 8603 Schwerzenbach, Switzerland; Centre for Stem Cell Biology, Department of Biomedical Science, University of Sheffield, Sheffield S10 2TN, United Kingdom

## Abstract

In contrast to conventional human pluripotent stem cells (hPSC) that are related to post-implantation embryo stages, naïve hPSC exhibit features of pre-implantation epiblast. Naïve hPSC are established by resetting conventional hPSC, or are derived from dissociated embryo inner cell masses. Here we investigate conditions for transgene-free reprogramming of human somatic cells to naïve pluripotency. We find that tankyrase inhibition promotes RNA-mediated induction of naïve pluripotency. We demonstrate application to independent human fibroblast cultures and endothelial progenitor cells. We show that induced naïve hPSC can be clonally expanded with a diploid karyotype and undergo somatic lineage differentiation following formative transition. Induced naïve hPSC lines exhibit distinctive surface marker, transcriptome, and methylome properties of naïve epiblast identity. This system for efficient, facile, and reliable induction of transgene free naïve hPSC offers a robust platform, both for delineation of human reprogramming trajectories and for evaluating the attributes of isogenic naïve versus conventional hPSC.

## INTRODUCTION

Human pluripotent stem cells (hPSC) provide a potent resource for fundamental research into early human development and in addition hold great promise for biomedical applications. hPSC have been derived by culture of explanted human embryo inner cell masses (ICM) (O’Leary et al., 2012; Thomson et al., 1998), and by reprogramming of somatic cells (Takahashi et al., 2007; Yu et al., 2007). The precise relationship between conventional hPSC and in vivo epiblast development is uncertain, but they have diverged from ICM (O’Leary et al., 2012; Yan et al., 2013) and appear to represent a post-implantation stage approaching gastrulation (Nakamura et al., 2016). Consequently these cells are often described as primed (Nichols and Smith, 2009; Rossant and Tam, 2017). A second type of hPSC has been isolated more recently using alternative culture conditions based on inhibition of the ERK pathway (Takashima et al., 2014; Theunissen et al., 2014). These cells are termed naïve because they show similarities to the pre-implantation epiblast (Guo et al., 2016; Stirparo et al., 2018; Theunissen et al., 2016) and may be analogous to the archetypal embryonic stem cells established in mouse (Nichols and Smith, 2012; Smith, 2001). Naïve hPSC are obtained by resetting the status of conventional hPSC using transgenes (Takashima et al., 2014) or by culture manipulation (Guo et al., 2017; Theunissen et al., 2014). Naïve cell lines can also be established directly from embryos after dissociation of the ICM (Guo et al., 2016).

Somatic cell reprogramming directed by ectopic transcription factors generates induced pluripotency (Takahashi and Yamanaka, 2006). The canonical Yamanaka reprogramming factors yield induced pluripotent stem cells (iPSC) that in mouse are naïve, but in human are primed (Okita et al., 2007; Silva et al., 2008; Takahashi et al., 2007). This difference may be determined by the appropriateness of the culture environment for capture of naïve versus primed states, respectively. Indeed, mouse primed iPSC can be obtained using medium containing FGF and activin (Han et al., 2011), similar to culture conditions for propagation of conventional hPSC (Vallier et al., 2005). Induction of naïve pluripotency is relatively robust in the mouse system and is increasingly well characterized at the molecular level (Guo et al., 2019; Schiebinger et al., 2019; Stadhouders et al., 2018). Reprogramming of human fibroblasts to naïve iPSC has only recently been reported, however, and appears variable and inefficient (Kilens et al., 2018; Liu et al., 2017). The methods entailed protracted reprogramming factor expression from viral or episomal vectors and the iPSC frequently exhibited persisting transgenes. Moreover the reprogrammed cells obtained were heterogeneous with poorly characterized differentiation behavior. Very recently, reprogramming to the human naïve state was achieved using modified mRNA vectors applied in a microfluidic apparatus (Giulitti et al., 2019). In that study the authors report that serial transfection with chemically modified mRNAs over at least 7 days within a microfluidic chamber is important for induction of naïve cells. Thus human naïve reprogramming contrasts with findings in mouse in which naïve iPSC are readily obtained by multiple methods requiring only short-term exposure to reprogramming factors in standard tissue culture conditions.

Here we sought to determine whether human naïve iPSC could be produced directly from somatic cells in bulk culture with simplicity and efficiency comparable to the generation of mouse iPSC. Integration and/or persisting expression of reprogramming factor transgenes is undesirable in principle, and specifically may perturb the naïve PSC state or subsequent differentiation. We therefore focused on producing transgene-free naïve hPSC by transient delivery of non-modified RNAs (Poleganov et al., 2015).

## RESULTS

### RNA-mediated induction of naïve pluripotency is facilitated by tankyrase inhibition

RNA-directed reprogramming has previously been used to generate conventional human iPSC (Poleganov et al., 2015). We reasoned that the same system may induce naïve pluripotency under the appropriate culture conditions. We adopted the combination of mRNAs encoding six reprogramming factors, OCT4, SOX2, KLF4, c-MYC, NANOG and LIN28 (OSKMNL), augmented with microRNAs 302 and 367, and supplemented with Vaccinia virus immune evasion factors E3, K3, and B18R mRNAs to suppress the interferon response. Naïve hPSC were originally established and propagated in medium containing the MEK1/2 inhibitor PD0325901 (P), the glycogen synthase kinase-3 (GSK3) inhibitor CH99021 (CH), the atypical protein kinase C inhibitor Gö6938 (Gö or G) and the cytokine leukemia inhibitory factor (LIF), collectively termed t2iLGö (Guo et al., 2016; Takashima et al., 2014). More recently, however, we have found that the tankyrase inhibitor and Wnt pathway antagonist XAV939 (XAV) enhances transgene-free resetting of conventional PSC to naïve status (Bredenkamp et al., 2019; Guo et al., 2017). We therefore examined the respective effects of CH and XAV during RNA-mediated reprogramming.

We plated 10,000 human diploid fibroblasts (HDFs) on Geltrex coated 4-well tissue culture plates and after overnight incubation carried out transfections with the RNA cocktail for four consecutive days (Fig 1A). Cells were then cultured in medium containing FGF2 for two days before exchange to naïve reprogramming media. The naïve media each contained PD0325901 (1µM), Gö6938 (2µM) and human LIF (10ng/ml), plus the Rho-associated coiled-coil kinase (ROCK) inhibitor Y27632 (1µM). To this base medium, termed PGL, we added either CH (1µM), as in the original t2iLGö naïve hPSC culture formulation (Takashima et al., 2014), or XAV (2µM), constituting PXGL.

**Figure 1.**
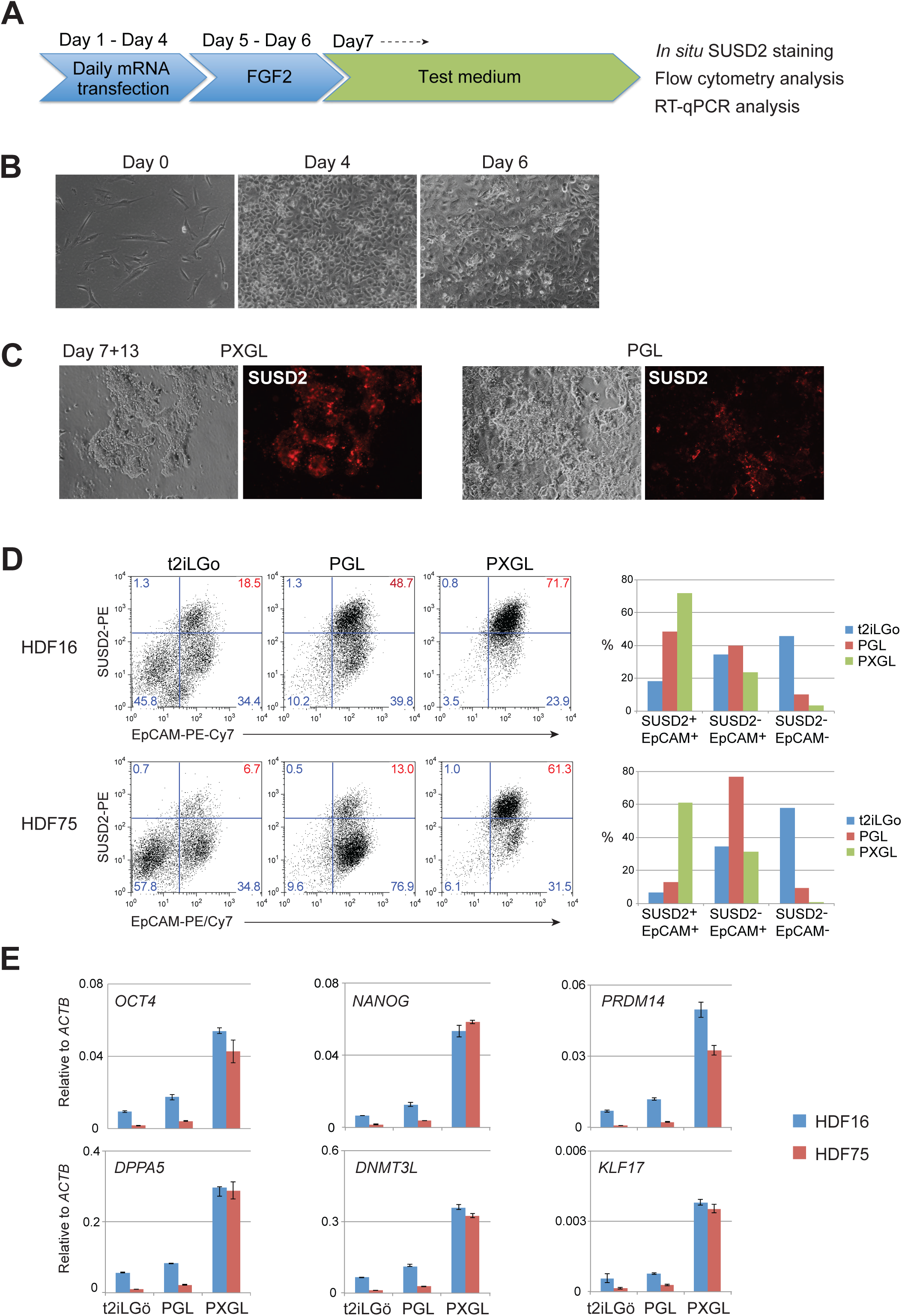
Tankyrase inhibition enhances naive reprogramming by RNA. 1A. Schematic of reprogramming protocol. 1B. Morphology during initial reprogramming in medium with FGF2. 1C. Morphology in naïve capture medium, PGL or PXGL. See also Figure S1A. 1D. Flow cytometry analysis of EpCAM and SUSD2 expression after 12 days in PGL with CH or XAV. Scatter plots on left, histograms on right. 1E. RT-qPCR analysis of pluripotency markers after 12 days in PGL based medium.

Fibroblasts grew to a near confluent layer of cells by the end of RNA transfection, and patches of cells undergoing mesenchymal epithelial transition became apparent from day 6 (Figure 1B, S1A). Following transfer to PGL-based naïve media we observed compact colonies of cells with smooth boundaries after a further 10 days (Figure S1A). We noted that addition of XAV resulted in markedly more of these colonies and a corresponding reduction in alternative cell morphologies.

Sushi Domain Containing 2 (SUSD2) is a cell surface protein highly expressed by human pre-implantation epiblast cells and naïve hPSC (Bredenkamp et al., 2019). By in situ live staining we detected expression of SUSD2 by the majority of compact colonies in reprogramming cultures in the presence of XAV (Figure 1C, S1A). We quantified the effect of XAV or CH by flow cytometry using SUSD2 together with the pan-epithelial marker EpCAM. The proportion of SUSD2^+^EpCAM^+^ cells was substantially higher in the presence of XAV than in PGL. Conversely, CH reduced the number of SUSD2+EPCAM+ cells (Figure 1D). Consistent with SUSD2 analysis, cultures reprogrammed in the presence of XAV showed substantially higher expression of core pluripotency factors and of naive markers assayed by RT-qPCR (Figure 1E).

### Reproducibility of reprogramming to naïve status

Somatic cell reprogramming can vary between cell lines. To evaluate reproducibility of RNA-directed reprogramming to a naïve phenotype we applied the protocol using PXGL to cultures of two adult primary dermal fibroblasts (HDF16, HDF75) and one newborn foreskin fibroblast (BJ). The experiments were repeated at different passages and we tested three different batches of RNA cocktail. In all cases we obtained SUSD2 positive colonies. SUSD2 live staining after 12-14 days in PXGL typically revealed several 100 stained colonies per well of a 4-well plate (Figure 2A, S1B). To substantiate the character of these colonies we carried out immunostaining for diagnostic transcription factors. KLF17 is a transcription factor expressed in the early human embryo and in naïve PSC but entirely absent from conventional PSC (Blakeley et al., 2015; Guo et al., 2016), and NANOG is a critical pluripotency factor expressed in both naïve and conventional hPSC. We detected co-expression of KLF17 and NANOG proteins in the majority of reprogrammed colonies in PXGL (Figure 2B, S1C).

**Figure 2.**
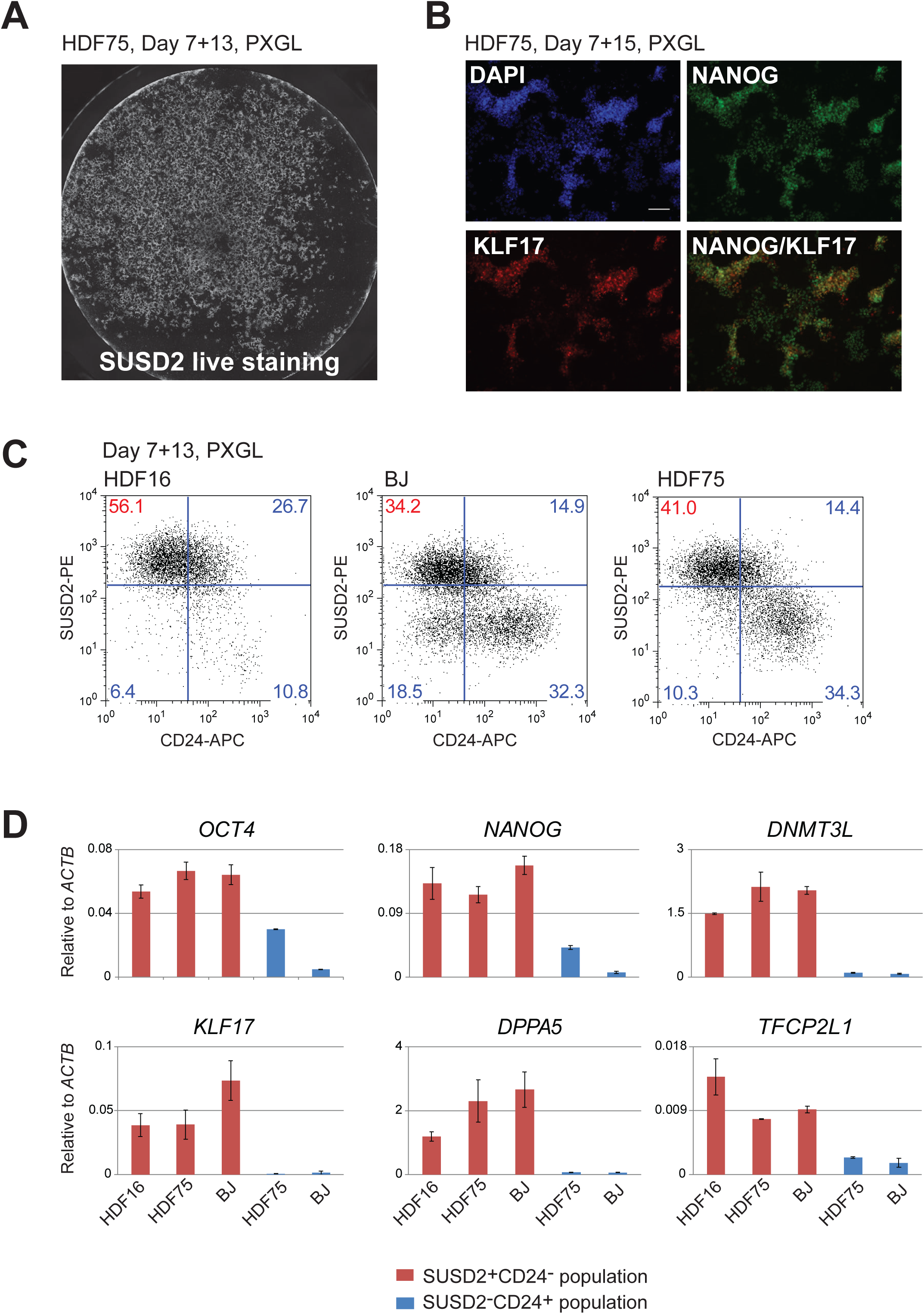
Reproducibility of reprogramming in PXGL. 2A. Well of HDF75 reprogramming culture after 13 days in PXGL, stained in situ with SUSD2-PE antibody. See also figure S1B. 2B. Immunostaining for KLF17 and NANOG after 15 days in PXGL. 2C. Flow cytometry analysis SUSD2 and CD24 expression at day 13 in PXGL for different fibroblast lines. 2D. Marker analysis by RT-qPCR of isolated SUSD2 positive and negative populations. Error bars are SD from 2 technical replicates.

Human naïve and conventional PSC are distinguished by exclusive expression of either SUSD2 or CD24 surface markers respectively (Bredenkamp et al., 2019). We quantified naïve reprogramming for HDF16, HDF75 and BJ cultures based on presence of SUSD2 and absence of CD24 after 14 days in PXGL (Figure 2C). For HDF16, the majority of the culture (56%) comprised SUSD2^+^CD24^-^ cells. BJ and HDF75 cultures were more mixed at this stage. In addition to SUSD2 positive cells, a distinct SUSD2^-^CD24^+^ population was also present. We purified these two populations and subjected them to RT-qPCR analysis. SUSD2^+^ cells express naïve markers KLF17, KLF4, TFCP2L1, DPPA5 and DNMT3L, while the CD24^+^SUSD2^-^ population express general pluripotency markers OCT4 and NANOG at low levels but lack naïve hallmarks (Figure 2D).

We investigated reprogramming of an alternative somatic cell type, peripheral blood-derived endothelial progenitor cells, EPC (Ormiston et al., 2015). EPC require more prolonged RNA transfection over 8 days (Poleganov et al., 2015) and incur considerable cell death. Surviving cells were transferred to PXGL and after three weeks we observed occasional patches of compact epithelial cells. We detected SUSD2 in 5% of the population, and observed expression of KLF17 and NANOG proteins (Figure S2).

### Expansion of naïve iPSC generated by RNA-mediated reprogramming

After 14 days in PXGL for HDF and 21 days for EPC, we bulk passaged cultures via dissociation with accutase and replated onto feeder layers of mouse embryo fibroblasts (MEF) in PXGL plus ROCK inhibitor. Dome-shaped, refractile, colonies formed on MEF (Figure 3A). After 2 passages, HDF16- and BJ-derived cultures consisted of more than 90% SUSD2 positive cells (Figure 3B). HDF75- and EPC-derived cultures remained heterogeneous, however. In these cases we used flow cytometry to purify the SUSD2^+^CD24^-^ population. Thereafter we found that cells could readily be maintained with relatively homogeneous naïve colony morphology and SUSD2 expression (Figure 3C). Cultures were passaged every 4-5 days at a 1:3 or 1:5 split ratio for at least 6 weeks (> 10 passages). Expanded cultures display naïve transcription factor proteins KLF17, KLF4, and TFCP2L1 (Figure 3D). RT-qPCR analysis showed expression of naïve markers at comparable levels to naïve HNES cells (Guo et al., 2016) derived from dissociated human ICM (Figure 3E).

**Figure 3.**
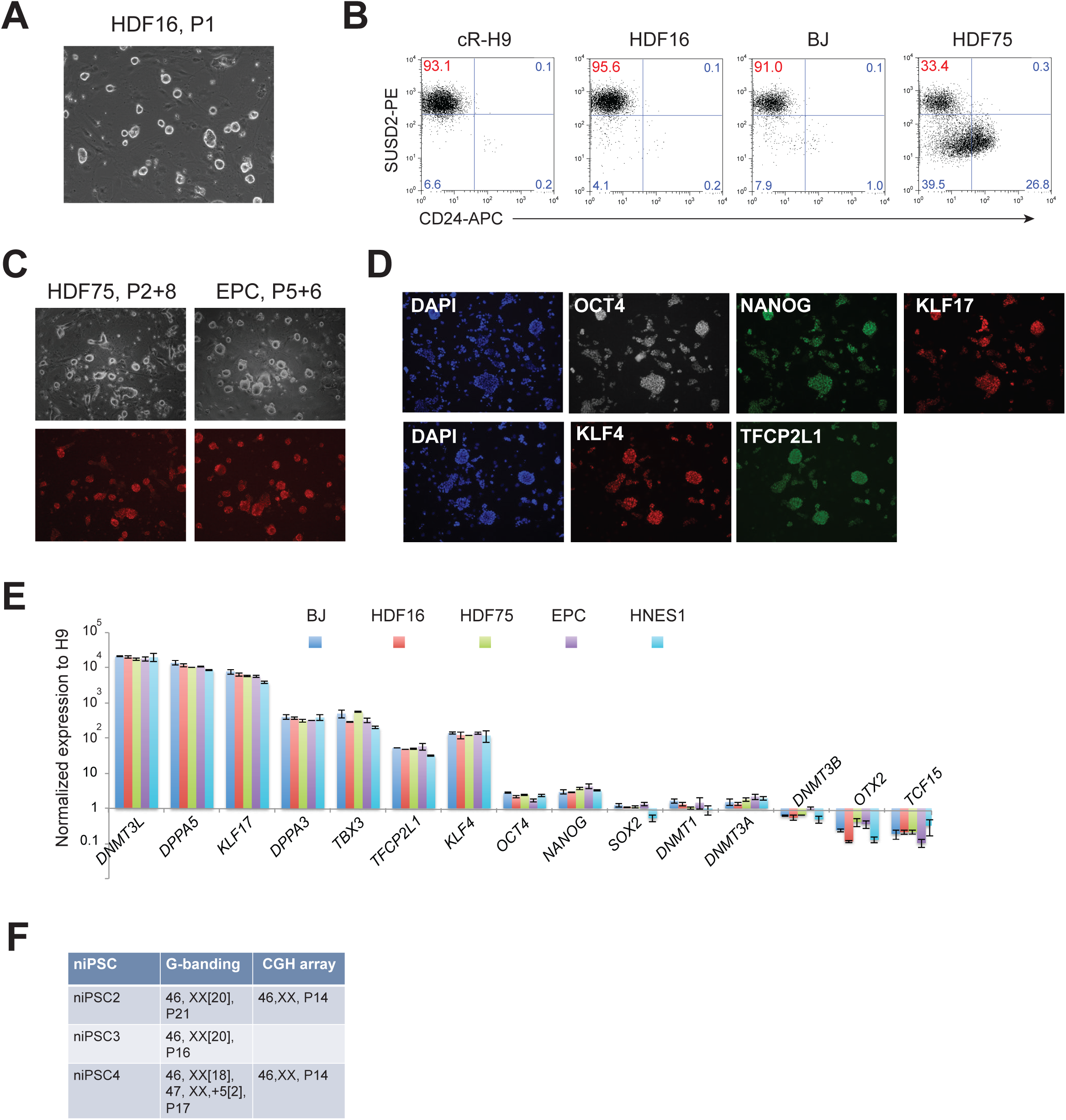
Expansion and characterization of naïve iPSCs. 3A. Morphology of naïve iPSC culture on MEF at passage 1 after reprogramming. 3B. Flow cytometry analysis of SUSD2 and CD24 expression in HDF16, HDF75 and BJ derived naïve iPSC cultures at passage 2. 3C. SUSD staining of naïve iPSC cultures of indicated origin after sorting and subsequent passaging (P). 3D. Immunostaining for naïve markers in expanded naïve iPSCs (BJ derived). 3E. RT-qPCR analysis of marker expression in expanded naïve iPSCs of indicated origins and in embryo-derived naïve HNES1 cells. Data are normalized to expression in conventional H9 cells. Error bars are SD from 2 technical replicates. 3F. Chromosome analysis of naïve iPSCs by G-banding and CGH array at indicated passage numbers (P). See also Figure S3B.

We also investigated expansion of individual colonies from the primary reprogramming well. We manually picked 8 colonies from HDF75 cultures after 14 days in PXGL. Colonies were dissociated with accutase and plated in PXGL plus ROCK inhibitor on MEF in a 96-well plate. Six colonies were expanded into stable naïve iPSC cultures that maintained naïve marker gene expression (Figure S3A).

We previously noted incidences of polyploidy in naive cells cultured in t2iLGö medium (Guo et al., 2016). We therefore monitored DNA content in the expanded naïve iPSC colonies by propidium iodide staining and flow cytometry analysis. One clone, niPSC1, contained a fraction of hyperdiploid cells at passage 5 but the other five clones remained diploid at passage 10 (Figure S3B). We performed G-banding karyotype analysis on three diploid clones, niPSC2, niPSC3 and niPSC4, after further expansion. niPSC2 and niPSC3, exhibited a normal 46, XX diploid karyotype at passages 20 and 16 respectively (Figure 3F). The third clone, niPSC4, was predominantly diploid but with a subpopulation (10%) of cells showing trisomy for chromosome 5 at passage 17. Array CGH on niPSC2 and niPSC4 at passage 14 confirmed a 46XX chromosome complement and did not detect any large DNA copy number variations. Therefore, human naïve iPSC can be expanded in PXGL with a relatively stable genome.

### Somatic lineage differentiation of naïve iPSC

Naïve PSCs represent pre-implantation epiblast and are not directly competent for somatic lineage induction (Rostovskaya et al., 2019; Smith, 2017). Formative transition of human naïve PSC can be achieved by transfer from to N2B27 medium supplemented with XAV only, a process termed capacitation (Rostovskaya et al., 2019). We examined differentiation potential of niPSC2 and niPSC4 following 13 days capacitation. Capacitated HNES1 and RNA reprogrammed isogenic primed iPSC were included for comparison. Both naïve iPSC clones differentiated efficiently to definitive endoderm, neuroectoderm andparaxial mesoderm upon directed lineage induction. For definitive endoderm, flow cytometry analysis quantified co-expression of SOX17 and CXCR4 in nearly 80% of cells after three days (Figure 4A). For each of the lineages, RT-qPCR and immunostaining detected expression of representative markers (Figure 4B-G, S4A-C).

**Figure 4.**
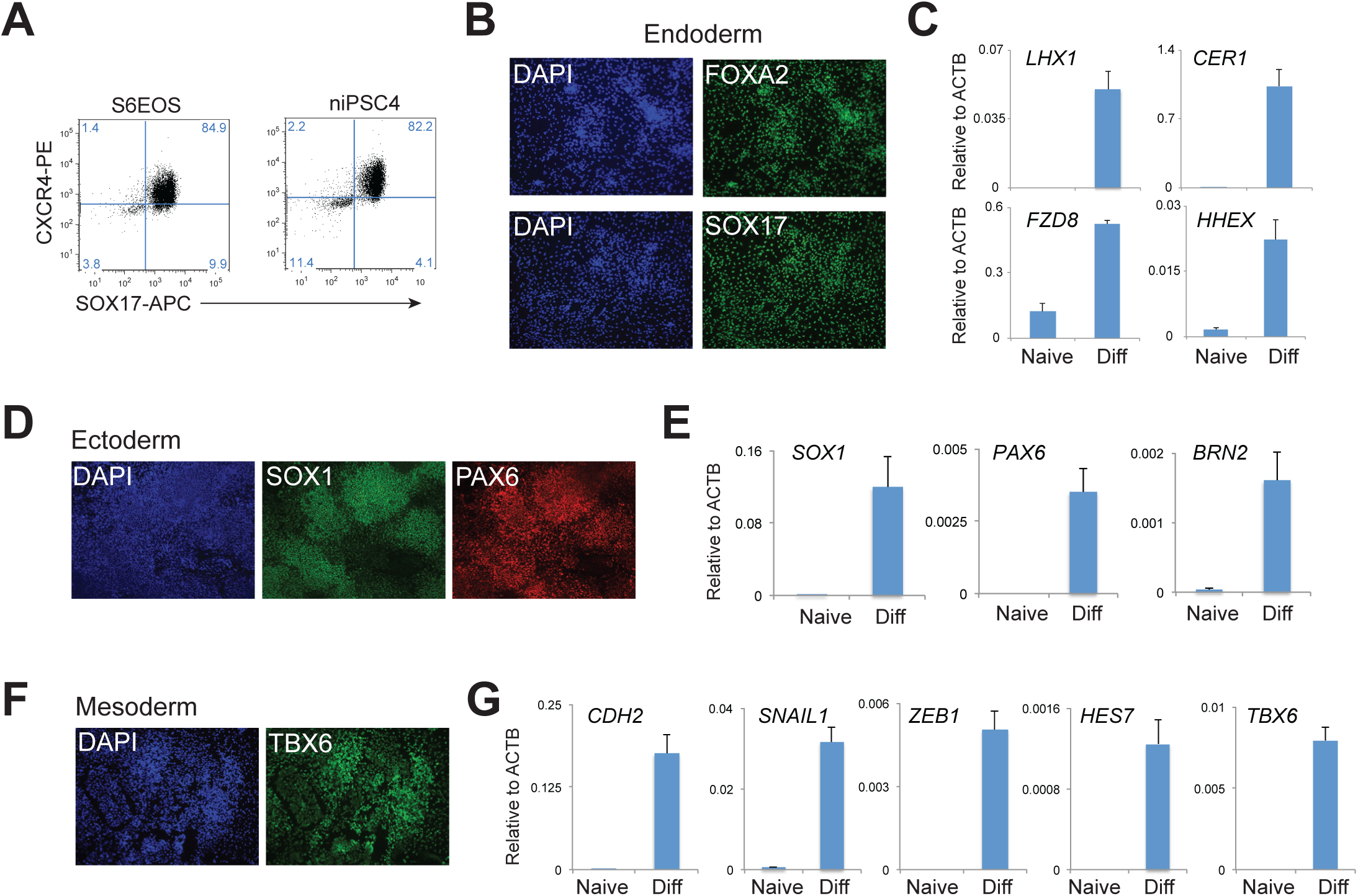
Differentiation of capacitated naïve iPSCs. 4A. Flow cytometry analysis of SOX17 and CXCR4 expression after 3 days definitive endoderm induction. 4B. Immunostaining for FOXA2 and SOX17 after 3 days definitive endoderm induction. 4C. RT-qPCR analysis of definitive endoderm markers after 3 days induction. 4D. Immunostaining for neuroectoderm markers SOX1 and PAX6. 4E. RT-qPCR analysis of neuroectoderm marker expression. 4F. Immunostaining for mesoderm marker TBX6. 4G. RT-qPCR analysis of paraxial mesoderm markers.

### Global transcriptome and DNA methylome features of naïve human iPSC

We carried out RNA-seq on niPSC2, niPSC4 and HNES1 passaged in PXGL on either geltrex or laminin to remove MEF. Two primed iPSC cultures generated by RNA-mediated reprogramming were examined in parallel. We applied quadratic programming (DeconRNAseq) to assess quantitatively the similarity between the PSC cultures and human pre-implantation development based on all expressed protein coding genes (Gong and Szustakowski, 2013; Stirparo et al., 2018). HNES1, niPS-C2 and niPS-C4 have a median epiblast fraction of identity of 0.8, 0.81, and 0.78 respectively (Figure 5A). These values indicate very high resemblance to pre-impantation epiblast compared to other stages (zygote, 4-cell, 8-cell, compacted morula, early ICM, primitive endoderm). In contrast the primed iPSC show less than 50% fraction of identity to epiblast.

**Figure 5.**
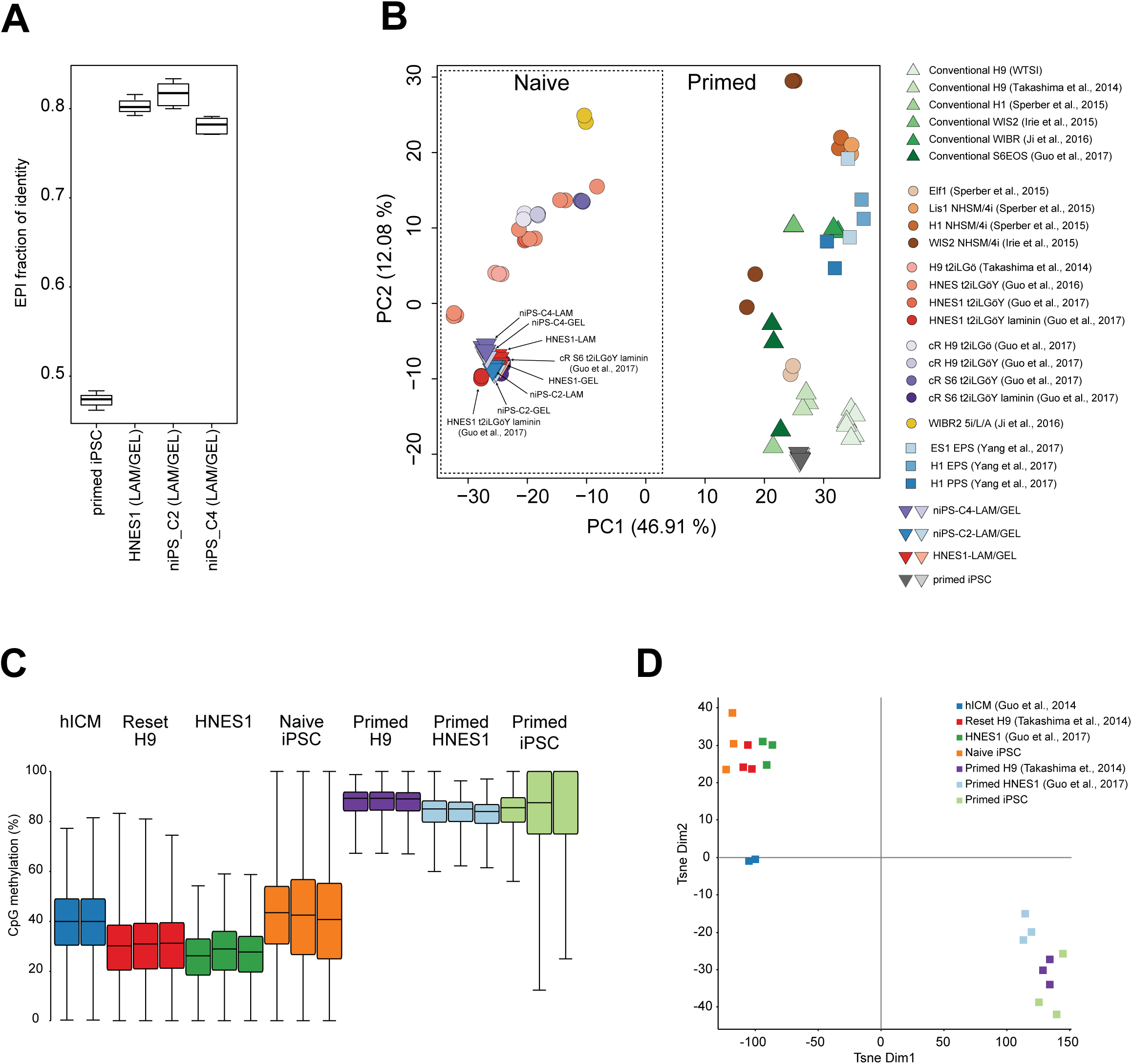
Global molecular analyses of naïve iPSCs. 5A. Fraction of identity with human pre-implantation epiblast for primed iPSC, HNES embryo-derived naïve stem cells, and naïve iPSCs. 5B. Principal component analysis for variable expressed genes. 5C. Box plots showing the global distribution of CpG methylation levels from pooled replicates of the indicated samples compared with published datasets (Guo et al., 2017; Guo et al., 2014; Takashima et al., 2014). iPSC samples are from independent experiments. Methylation was quantitated over 20 kb genomic tiles. 5D. t-SNE plot showing the distribution and clustering of the analyzed datasets. Methylation was quantitated over 20 kb genomic tiles.

We then compared these samples with previously analysed hPSC samples. Dimensionality reduction by principle component analysis (PCA) highlights that the naïve iPSC clones are very closely related to one another and to HNES1 cells cultured in PXGL, and also to naïve PSC cultured in a previous study in t2iLGö on laminin (Guo et al., 2017) (Figure 5B). Naïve PSC cultures on MEF in t2iLGö (Guo et al., 2017; Guo et al., 2016; Takashima et al., 2014) or 5iLA (Theunissen et al., 2014) are more dispersed in the PCA but reside in the same major cluster that is unambiguously separated on PC1 from various primed-type hPSC cultures.

Naïve hPSCs have been found to be globally hypomethylated (Takashima et al., 2014; Theunissen et al., 2016), in common with mouse and human ICM cells (Guo et al., 2014; Lee et al., 2014; Smith et al., 2014). To evaluate genome methylation in naive iPSCs, we performed whole-genome bisulfite sequencing. Methylation profiles for naïve and primed iPSC generated by RNA reprogramming were compared with published datasets for primed hPSCs, human ICM cells, transgene reset PSCs and HNES1 cells (Guo et al., 2016; Guo et al., 2014; Takashima et al., 2014). The primed iPSCs showed high levels of DNA methylation (85-95%), comparable with other primed hPSC samples generated in this study. In contrast, naïve iPSCs were globally hypomethylated to levels comparable to ICM cells but slightly higher than long-term cultures of transgene reset or embryo derived hPSCs (Figure 5A). Using t-Distributed Stochastic Neighbor Embedding (t-SNE) analysis (van der Maaten and Hinton, 2008), we also found that methylation profiles of naïve and primed PSC cultures clustered apart, with naïve cultures adjacent to ICM samples (Figure 5B).

Since we had previously shown that the genome of naïve PSCs is not completely hypomethylated, but exhibits a small number of regions that gain methylation compared to primed PSCs (Guo et al., 2017), we asked whether the naïve iPSC cultures displayed similar characteristics. We defined genomic regions (blue) which showed >10% hypermethylation between reset and primed H9-NK2 PSCs (Takashima et al., 2014) and <30% methylation in primed conditions and examined their methylation state in the current datasets (Figure S5A). We found that a substantial number of these regions were also hypermethylated in naïve iPSCs, indicating that this may be a general feature of naïve stem cells.

We also assessed the methylation status of imprinted regions in the different iPSC cultures. As observed previously (Guo et al., 2017; Pastor et al., 2016), naïve conditions failed to faithfully preserve imprinted methylation, although a significant number of imprints also appeared to be eroded in primed iPSC cultures (Figure S5B).

Overall, these analyses establish that human naïve iPSCs generated by RNA-directed reprogramming are related to human pre-implantation epiblast and are essentially indistinguishable globally from naïve PSC derived from human ICM or generated by resetting of conventional hPSC.

## DISCUSSION

The findings in this study establish that human somatic cells can be reprogrammed efficiently to the naïve PSC state by transient delivery of reprogramming factors using RNA transfection. Thereafter, naïve cells can reliably be expanded into stable diploid cell lines, either as bulk populations, by sorting for SUSD2 expression, or by picking individual colonies. Resulting naïve iPSC lines exhibit a consistent marker phenotype that is in common with previously characterized naïve hPSC produced by resetting or derived from embryos. Following formative transition, naïve iPSC display competence for differentiation into somatic lineages. Both transcriptome and DNA methylome of naïve iPSC show high global correlation with embryo derived naïve HNES cells and a corresponding relatedness to epiblast cells in the human blastocyst.

Recent studies reported that human naïve iPSC can be generated by transgene-induced reprogramming but that the products may be heterogeneous and confounded by persisting transgenes (Kilens et al., 2018; Liu et al., 2017). Transgene-free naïve iPSC have also been produced using chemically modified RNAs, but the efficiency of this approach was reported to depend on cell confinement in a microfluidic chamber (Giulitti et al., 2019), which restricts general application. In contrast, our results demonstrate that reprogramming to the naïve state can be highly efficient using unmodified RNAs in standard cell culture conditions. For dermal fibroblasts, three or four daily transfections with mRNAs encoding OSKMNL reprogramming factors together with miRNAs 302 and 367 are sufficient to produce more than one hundred SUSD2+ naïve iPSC colonies starting from 10,000 cells in a single well of a 4-well plate. This result is qualitatively reproducible between three different human fibroblast cultures, although individual efficiency varies, as has been generally noted for human reprogramming. Of note, PXGL medium not only promotes establishment of naïve pluripotency but is also relatively selective against other cell types. Consequently the majority of non-s or incompletely reprogrammed cells die or growth arrest in these conditions, generally allowing naïve iPSC cultures to be established by bulk passaging without need for colony picking or cell sorting, although both can also be deployed. Occasionally we noticed high levels of cell death during RNA transfection, in which case limiting the transfection period to three days preserves viability and naïve colonies are still generated in recoverable numbers. In the case of EPC, sustained transfection is required and reprogramming efficiency is lower, as also noted for conventional iPSC generation (Poleganov et al., 2015), but sorting for SUSD2^+^CD24^-^ cells effectively purifies the naïve cell fraction and enables subsequent stable expansion.

We found that supplementation with XAV markedly improves the efficiency of reprogramming to the naïve state, in line with observations during resetting of conventional PSC (Guo et al., 2017). This may be a key difference from previous reports that found low efficiency of naive reprogramming using media that typically included the GSK3 inhibitor CH (Giulitti et al., 2019; Kilens et al., 2018; Liu et al., 2017). Our analysis shows that the presence of CH inhibits reprogramming to naïve status. CH has the opposite effect to XAV of stimulating rather than suppressing canonical WNT signaling. We surmise that blockade of WNT signaling reduces activation of gene expression that can derail reprogramming and/or destabilise naïve hPSC, as evidenced during resetting (Guo et al., 2017). Thus insulation from WNT signaling appears beneficial for stabilisation of naïve pluripotency during induction and expansion. This is in line with the general proposition that naïve PSC are sustained primarily by preventing differentiation (Martello and Smith, 2014), though differs in detail from the mouse ground state system (Ying et al., 2008). The species difference may largely be explained by the fact that human naïve PSC, and in vivo human naïve epiblast cells, show very low expression of TCF3 (*TCF7L1*) and do not express ESRRB (Rostovskaya et al., 2019; Takashima et al., 2014), the key components regulated by GSK3 inhibition in mouse ES cells(Martello et al., 2012; Wray et al., 2011). In general, we find that stem cell cultures in PXGL exhibit equivalent naïve features to cells in our original t2iLGö formulation (Takashima et al., 2014), but appear more robust and stable.

Relatively facile but reliable generation of naïve iPSC as described here will open up the fields of human reprogramming and naïve pluripotency for deeper investigation. In mouse it is well-established that somatic cell reprogramming converges on the naïve PSC phenotype unless specific culture conditions are applied to capture primed pluripotency (Han et al., 2011). In human, however, the same reprogramming factors as used in mouse routinely generate PSCs of the primed phenotype. Our findings substantiate the hypothesis that the final state of pluripotency obtained by molecular reprogramming is determined in human as in mouse by the culture environment. We speculate that reprogramming to the naïve state may be direct in the PXGL culture environment and not entail passage through a primed state. It will be of interest to examine this by determining the trajectories of RNA-mediated reprogramming to naïve or primed endpoints. High efficiency with limited duration of reprogramming factor expression makes the mRNA delivery system attractive for such studies applied to primary cells. Furthermore, as illustrated in the case of XAV, it is straightforward to combine small molecules with RNA reprogramming and screen for accelerated or enhanced reprogramming, which can readily be visualised and quantified using SUSD2 live staining or flow cytometry (Bredenkamp et al., 2019). Finally, the ability to generate naïve iPSC rapidly and reliably from somatic cells provides a platform for comprehensive evaluation of the consistency, genomic stability, differentiation propensity, and other attributes of naïve hPSC compared to isogenic conventional hPSC generated from the same donor.

## MATERIALS AND METHODS

### Human PSC culture

Naïve hPSC including, chemically reset (cR), embryo-derived (HNES1) and naïve iPSCs were propagated in N2B27 with PXGL [1 µM PD0325901 (P), 2 µM XAV (X), 2 µM Gö6983 (G) and 10 ng/ml human LIF (L),] on irradiated MEF feeders. ROCK inhibitor (Y-27632) and Geltrex (2μg per cm^2^ surface area; Thermo Fisher, A1413302, growth factor-reduced) were added to media during replating. Cells were cultured in 5% O_2_, 7% CO_2_ in a humidified incubator at 37°C and passaged by dissociation with Accutase (Thermo Fisher Scientific, A1110501) every 3-5 days. For capacitation, cells were passaged once without feeders in PXGL medium then exchanged into N2B27 containing 2µM XAV (Rostovskaya et al., 2019). Conventional hPSC cultures were propagated on Geltrex in Essential 8 (E8) medium made in-house (Chen et al., 2011) or AFX medium (N2B27 basal medium with 5ng/ml Activin A, 5ng/ml FGF2 and 2 µM XAV). Cell lines were maintained without antibiotics and confirmed free of mycoplasma contamination by periodic in-house PCR assay.

### Somatic cell culture

Adult human dermal fibroblasts (HDFa), HDFa16, HDFa75 (Thermo Fisher Scientific C0135C), and BJ foreskin fibroblast (ATCC® CRL-2522™) were cultured in DMEM high glucose (Sigma D5546) with FBS (10%, Sigma, F0804), L-glutamine (2mM, Invitrogen, 25030-024) and 2-mercaptoethanol (100μM, Sigma, M3148) on gelatin-coated plates. Peripheral blood-derived endothelial progenitor cells (EPC, C26b) were cultured as described (Ormiston et al., 2015) in endothelial cell basal medium (PromoCell, c-22210) supplemented with 10% FBS and cytokines without heparin.

### RNA Reprogramming

Reprogramming was performed using the *StemRNA 3rd Gen Reprogramming Kit* (Stemgent Code: 00-0076). A detailed protocol is provided in supplemental information. Briefly, fibroblasts were plated in culture medium with serum. The following day RNAs were delivered by Lipofectamine® RNAiMAX™ and transfection repeated daily for 3-4 days in medium supplemented with FGF2. From day 7 cultures were exchanged to naïve culture medium until naïve-type colonies formed.

### hPSC differentiation

Naïve hPSC capacitation and tri-lineage differentiation were performed as described (Rostovskaya et al., 2019). In brief, naïve PSCs were capacitated for more than 10 days to prepare them for lineage induction. Definitive endoderm was induced over three days: day 1 in CDM2 basal medium supplemented with 100 ng/ml, activin A, 100 nM PI-103,3 μM CHIR99021, 10 ng/ml FGF2, 3 ng/ml BMP4, 10 μg/ml heparin and followed by 2 days in CDM2 supplemented with 100 ng/ml activin A, 100 nM PI-103, 20 ng/ml FGF2, 250 nM LDN193189, 10 μg/ml heparin (Loh et al., 2014). Neuroectoderm was induced in N2B27 medium supplemented with 1μM A8301 and 500 nM LDN193189 for 10 days (Chambers et al., 2009). Differentiation to paraxial mesoderm was induced for 6 days in 3 μM CHIR99021 and 500 nM LDN193189, with addition of 20 ng/ml FGF2 from day 3-6 (Chal et al., 2015).

### Reverse transcription and real-time PCR

Total RNA was extracted using RNeasy Kit (Qiagen) and cDNA synthesized with SuperScript III reverse transcriptase (Thermo Fisher Scientific, 18080085) and oligo(dT) adapter primers. TaqMan assays and Universal Probe Library (UPL) probes (Roche Molecular Systems) were used to perform gene quantification.

### Immunostaining

Cells were fixed with 4% PFA for 10 min at room temperature and blocked/permeabilised in PBS with 0.1% Triton X-100, 3% Donkey serum for 30 min. Incubation with primary antibodies was overnight at 4°C. Wash in 0.1% Triton X-100 twice, 10 minute each time. Secondary antibodies were added for 1 h at room temperature. The following antibodies were used for immunostaining of pluripotency markers: NANOG (R&D Systems AF1997), OCT4 (Santa Cruz sc-5279), KLF4 (Santa Cruz sc-20691), KLF17 (Atlas Antibodies HPA024629), TFCP2L1 (R&D Systems AF5726). Antibodies for immunostaining of differentiation markers were: FOXA2 (R&D Systems AF2400), SOX17 (R&D Systems AF1924), SOX1 (R&D Systems AF3369), PAX6 (Merck Millipore AB2237), TBX6 (Abcam ab38883). For live staining, cells were incubated with conjugated SUSD2 clone W5C5 (SUSD2-PE, BioLegend 327406) in culture media for 30 min before washing and imaging.

### Flow cytometry

Flow cytometry analysis was carried out on a CyAn™ ADP (Beckman Coulter) or BD LSRFortessa instrument (BD Biosciences) with analysis using FlowJo software. For intracellular marker staining, cells were fixed with Fixation Buffer (00-8222-49, ThermoFisher Scientific) for 30min at +4°C, washed with Permeabilization Buffer (00-8333-56, ThermoFisher Scientific), and incubated with SOX17 antibody diluted with Permeabilization Buffer and 5% donkey serum (Sigma-Aldrich) for 1 hour at +4°C. Cell sorting was performed using a MoFlo high-speed instrument (Beckman Coulter). The following antibodies were used for flow cytometry: SUSD2-PE (BioLegend 327406), CD24-APC (eBioscience17-0247-42), EpCAM-PE/Cy7 (BioLegend 324221), Tra1-85-FITC (Miltenyi Biotec 130-107-106), CXCR4-PE (BD Pharmingen 555974) BD Pharmingen, SOX17-APC (R&D Systems IC1924A).

### Chromosome analysis

Metaphase spreads were prepared and G-banded by standard procedures, as described (Guo et al., 2017). CGH array analysis using the Agilent ISCA 8×60K v2 array was carried out at the Cytogenetics Laboratory, Cambridge University Hospitals.

### Transcriptome sequencing and data analysis

Naïve hPSCs were cultured on geltrex or laminin without MEF for three passages before harvesting for RNA. Total RNA was extracted from three biological replicates of each cell line using TRIzol/chloroform (Invitrogen) and RNA integrity assessed by Qubit measurement and RNA nanochip Bioanalyzer. Ribosomal RNA was depleted from 1 µg of total RNA using Ribozero (Illumina Kit). Sequencing libraries were prepared using the NEXTflex Rapid Directional RNA-Seq Kit. Sequencing was performed on the Illumina HiSeq4000 in paired end 125bp format.

Reads were aligned to human genome build GRCh38/hg38 with STAR (Dobin et al., 2013) using the human gene annotation from Ensembl release 87 (Yates et al., 2016). Alignments to gene loci were quantified with HTseq-count (Anders et al., 2014) based on annotation from Ensembl 87 and using option –m intersection-nonempty. Fractional identity between *in vitro* cultured cells and pre-implantation stages and was computed using R package DeconRNASeq (Gong and Szustakowski, 2013) and method as described (Stirparo et al., 2018). External datasets used for comparative analyses are detailed elsewhere (Guo et al., 2017; Stirparo et al., 2018). Principal component analyses were performed based on log_2_ FPKM values computed with the Bioconductor packages *DESeq2 (Love et al., 2014)* or *FactoMineR* (Lê et al., 2008) in addition to custom scripts.

### Whole genome bisulfite sequencing, mapping, and analysis

Post-bisulfite adaptor tagging (PBAT) libraries for whole-genome DNA methylation analysis were prepared from purified genomic DNA (Miura et al., 2012; Smallwood et al., 2014; von Meyenn et al., 2016). Paired-end sequencing was carried out on HiSeq2500 instruments (Illumina). Raw sequence reads were trimmed to remove poor quality reads and adapter contamination using Trim Galore (v0.4.1) (Babraham Bioinformatics). The remaining sequences were mapped using Bismark (v0.14.4) (Krueger and Andrews, 2011) to the human reference genome GRCh37 in paired-end mode as described (von Meyenn et al., 2016). CpG methylation calls were analysed using SeqMonk software (Babraham Bioinformatics). Global CpG methylation levels of pooled replicates were illustrated using box plots. The SeqMonk build-in tSNE analysis was used to generate tSNE plots of the various datasets. The genome was divided into consecutive 20 kb tiles and percentage methylation was calculated using the bisulfite feature methylation pipeline in SeqMonk. Scatter plots of methylation levels over 20 kb tiles were generated using R, highlighting hypermethylated DMRs. Annotations of human germline imprint control regions were obtained as described(Court et al., 2014). Pseudocolour heatmaps representing average methylation levels were generated using the R heatmap.2 function without further clustering, scaling or normalisation.

### Data availability

RNA-seq and WGBS data are deposited in GEO database for release upon publication.

## Acknowledgements

We thank Jing Liu for RNA-sequencing. Peter Humphreys and Darran Clements supported imaging studies. Amer Rana kindly provided EPC line, C26b. We thank Sarah Eminli-Meissner and ReproCell for providing RNA reprogramming reagents.

## Author Contributions

Conceptualization, G.G; Methodology, G.G.; Investigation, G.G., N.B., J.Y., J.C., D.B., R.D., M.R., A.S.; Formal analysis, G.G.S., F.vM.; Writing, A.S and G.G.; Supervision G.G. and A.S.

## Supplementary protocol

Reprogramming human dermal fibroblasts to naïve pluripotent stem cells

### Materials

*HDFa (human dermal fibroblast, adult) Irradiated mouse embryonic fibroblast (MEF)*

*StemRNA 3rd Gen Reprogramming Kit* (Stemgent Code: 00-0076)

*Lipofectamine® RNAiMAX*™ (Invitrogen)

*Geltrex* (growth factor-reduced, Thermo Fisher, A1413302)

### Culture media

*Fibroblast culture medium*

DMEM high glucose (Sigma D5546), FBS (10%, Sigma, F0804), L-glutamine (2mM, Invitrogen, 25030-024), 2-mercaptoethanol (100μM, Sigma, M3148)

*Modified E7 medium*

Home made E6 basal medium (Chen et al., 2011) supplemented with 10 nglml FGF2 (prepare in house)

*NutriStem™ XF/FF Culture Medium* (Stemgent 01-0005)

*Naïve hPSC medium, PXGL*

N2B27 medium supplemented with MEK inhibitor PD0325901 (1μM), Tankyrase inhibitor XAV939 (2 μM), aPKC inhibitor Gö6983 (2μM), human LIF (10ng/ml, prepare in house)), Rho-kinase inhibitor Y-27632 (10μM)

### Protocol

1. **Day 0**: Dissociate HDFs with TrypLE. Collect dissociated cells, pellet at 300g for 3 minutes, and resuspend in fibroblast culture. Count cells and plate cells at a density of 1×10^4^/cm^2^ on tissue culture plate pre-coated with Geltrex.
2. **Day 1**: Switch to modified E7 medium and perform mRNA transfection following recommend of StemRNA™-NM Reprogramming Kit protocol.
3. **Day2-Day4**: Repeat mRNA transfection daily. **Note**: If excessive cell death after mRNA transfection is observed, higher number of fibroblasts is recommended. Alternatively, reducing mRNA transfection to 3 days can also rescue the culture and normally sufficient naïve colonies (more than 20) can be generate.
4. **Day5-Day6:** Refresh culture with modified E7 medium. NutriStem™ can be used as an alternative medium. **Note**, by day 6 patches of cells with epithelial morphology should become apparent at this stage, suggesting reprogramming has been induced by mRNA cocktail.
5. **Day7**: Switch to human naïve culture medium, PXGL and continue culture for about two weeks. SUSD2 positive colonies should appear about 7-10 days after in PXGL medium. Rock inhibitor (Y-27632) may be added to PXGL medium during reprogramming. **Note**, PXGL medium switch can be done between Day 6-Day9. Switching medium after Day10 will significantly reduce reprogramming efficiency.

### Passage and culture of established naïve iPSC

Human naïve iPSC are routinely cultured on MEFs at a density of 1-2×10^5^ per 6 well. The optimum feeder density needs to be tested experimentally for each batch of feeders. The MEF plates are prepared 3-7 days before use to allow MEFs spread evenly.

*Passaging naïve cells:*

1. Dissociate culture with Accutase or TrypLE Express (this will need about 5-10 minutes).
2. Pellet cells at 300g for 3 min. Aspirate and re-suspend cells in PXGL with Y-27632 (PXGLY).
3. Aliquot cells to new feeder plate with PXGLY. We recommend adding Geltrex (2ng per cm^2^) to cells at the time of passaging to aid attachment.
4. The next day, top up culture with fresh PXGL medium. Subsequently, half-medium changes daily until passaging. This normally takes 4-5 days culture in PXGL medium.

## Supplemental Figure Legends

**Figure S1.**
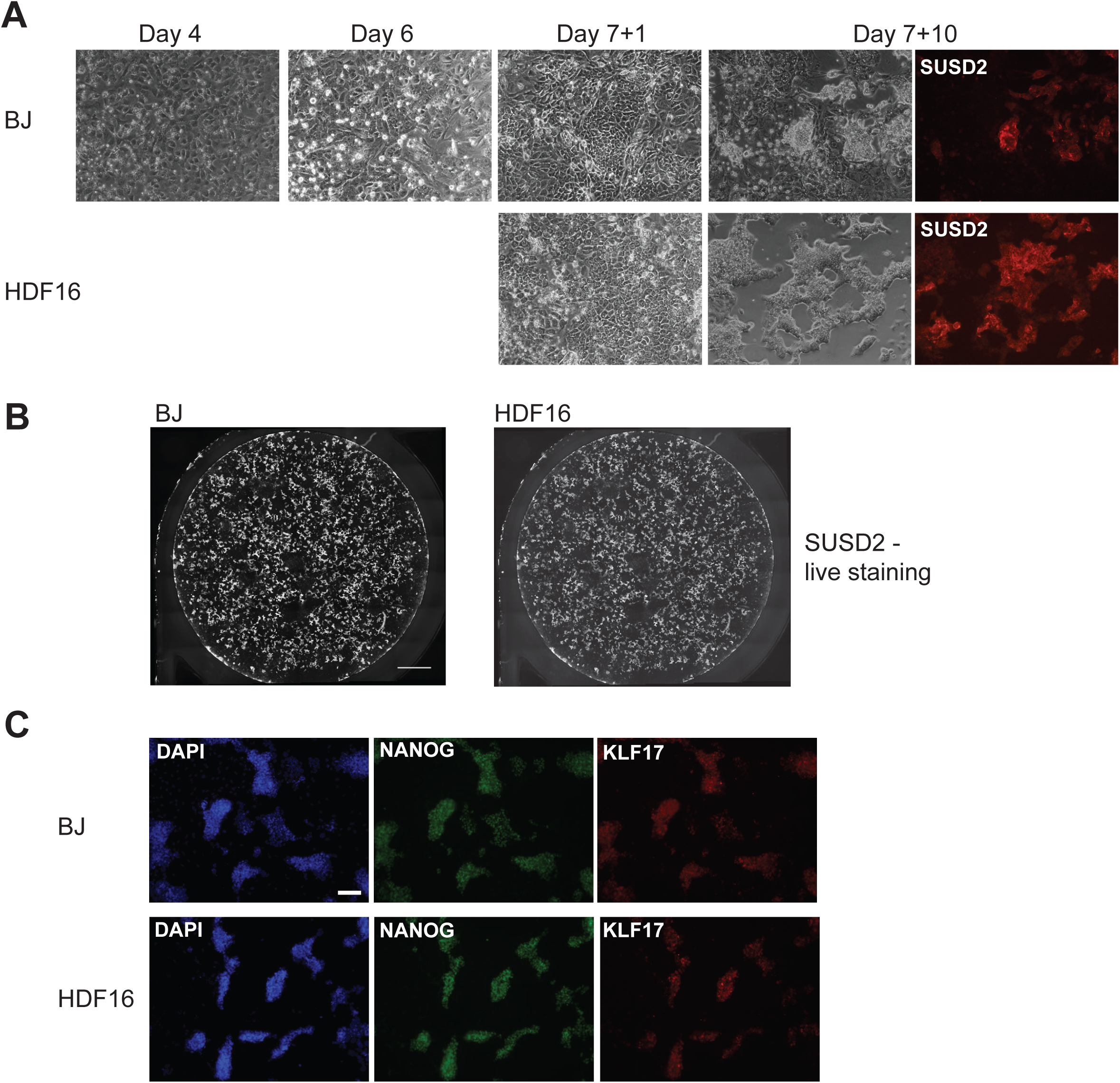
Reprograming of BJ and HDF16 cells. S1A. Morphology during reprogramming S1B. Wells of reprogramming cultures after 13 days in PXGL, stained in situ with SUSD2-PE antibody. S1C.Immunostaining for KLF17 and NANOG after 15 days in PXGL.

**Figure S2.**
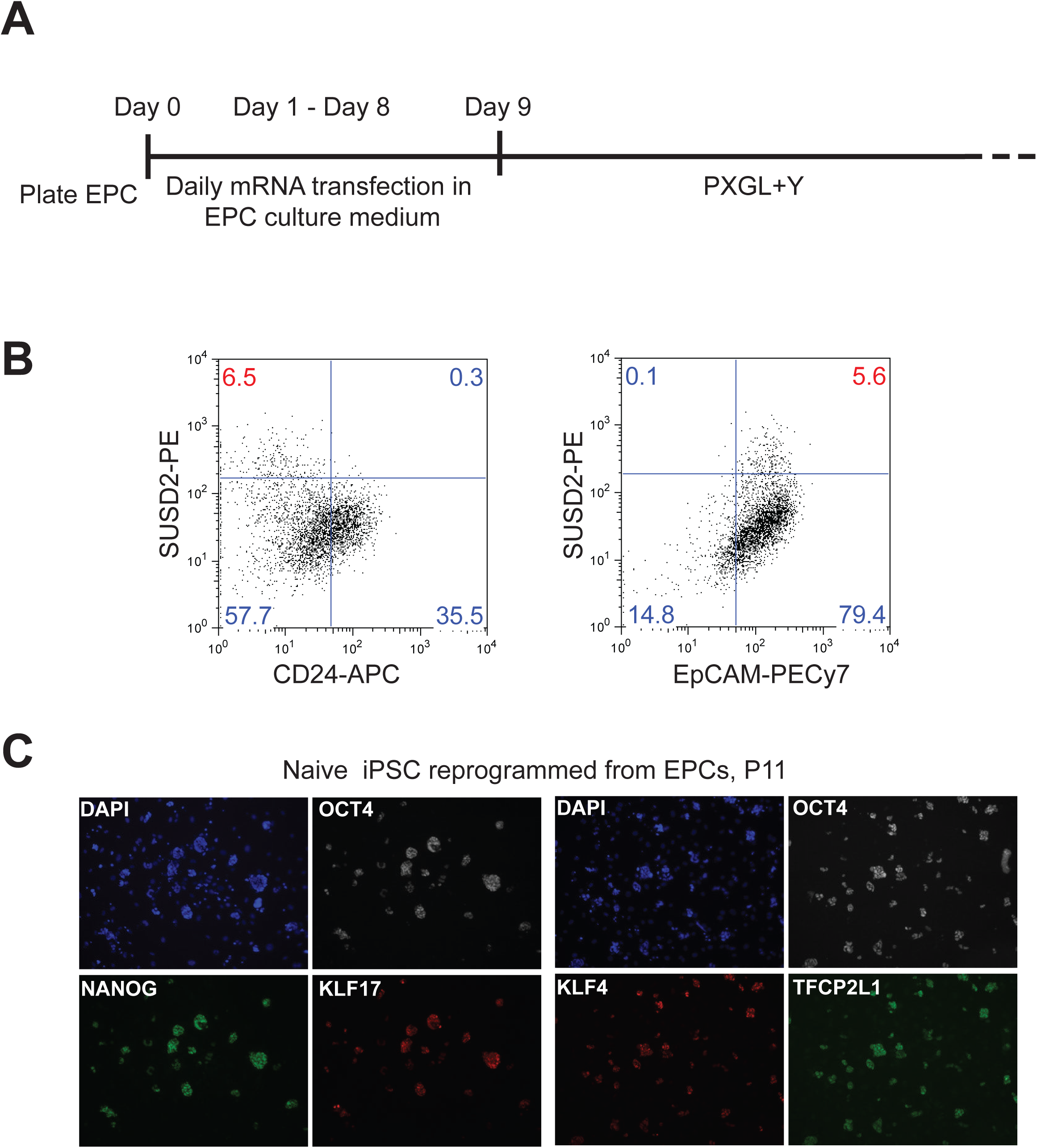
EPC reprogramming. S2A. Schematic of EPC reprogramming protocol S2B. Flow cytometry analysis of SUSD2, CD24 and EpCAM expression after three weeks in PXGL. S2C. Immunostaining of pluripotency markers in expanded EPC-derived naive iPSCs.

**Figure S3.**
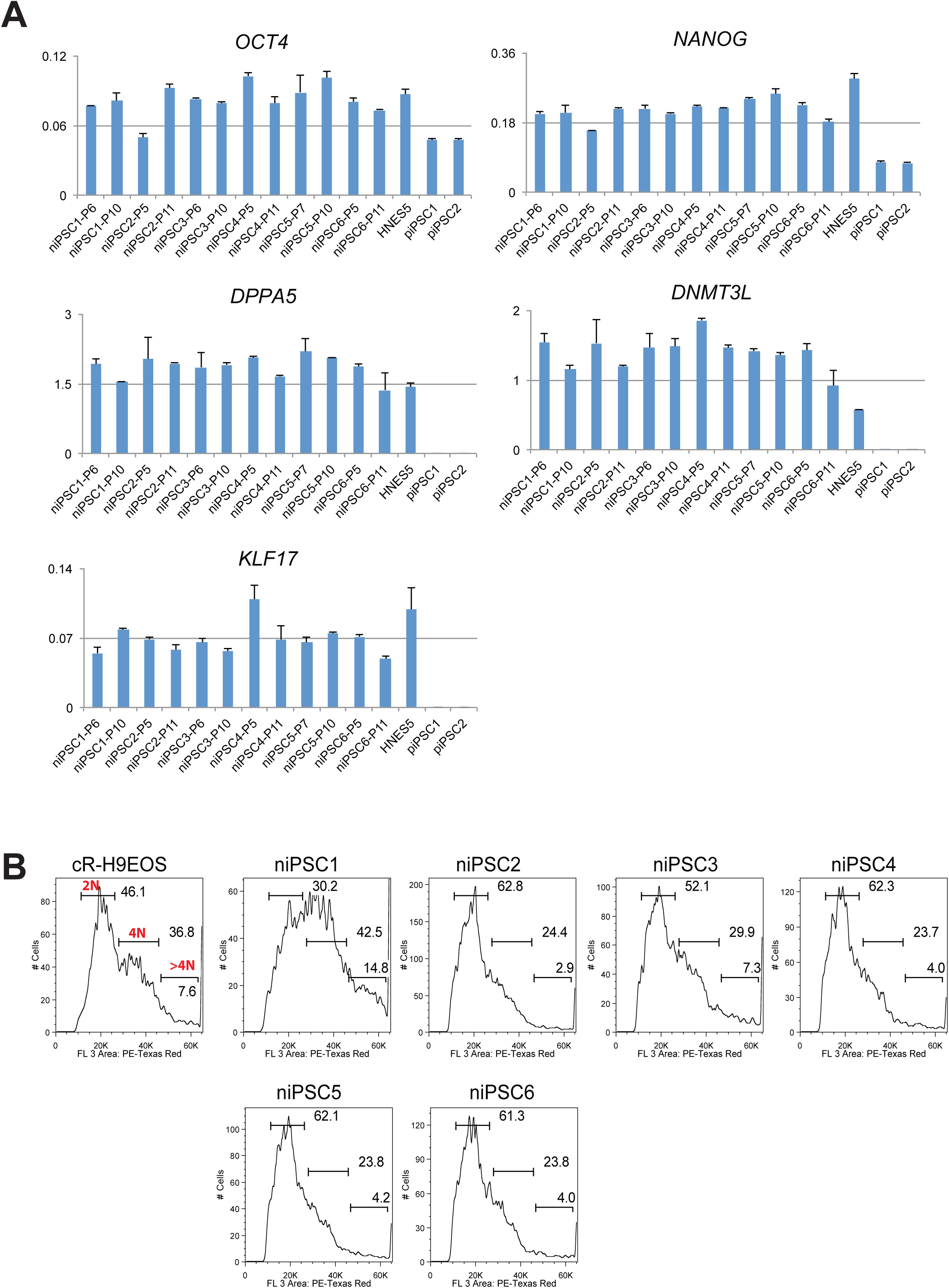
Clonal expansion of naïve iPSCs. S3A. RT-qPCR analysis of pluripotency markers in six expanded naïve iPSC clones at indicated passages. Two isogenic conventional iPSC clones (piPSC1, pIPSC2) generated in parallel and embryo derived HNES5 cells are included for comparison. Error bars are SD from two technical replicates. S3B. DNA content analysis from flow cytometry profiles of cells stained with propidium iodide. Diploid genome population is labeled as 2N, 4N indicates cells in G2 and/or tetraploid, hyperpolypoid is >4N.

**Figure S4.**
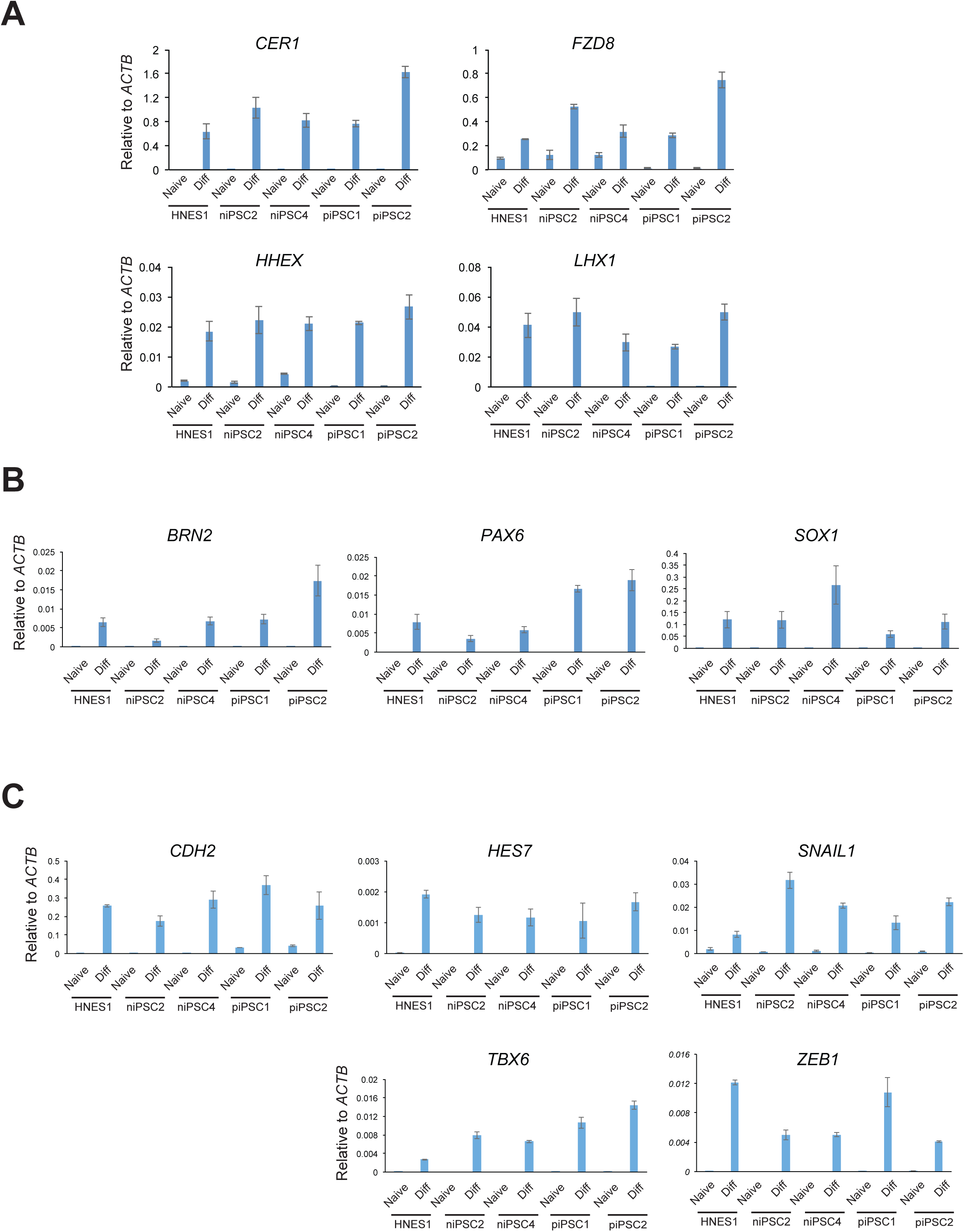
RT-qPCR analysis of lineage induction. S4A. Definitive endoderm markers S4B. Neuroectoderm markers S4C. Paraxial mesoderm markers Error bars are SD of technical duplicates.

**Figure S5.**
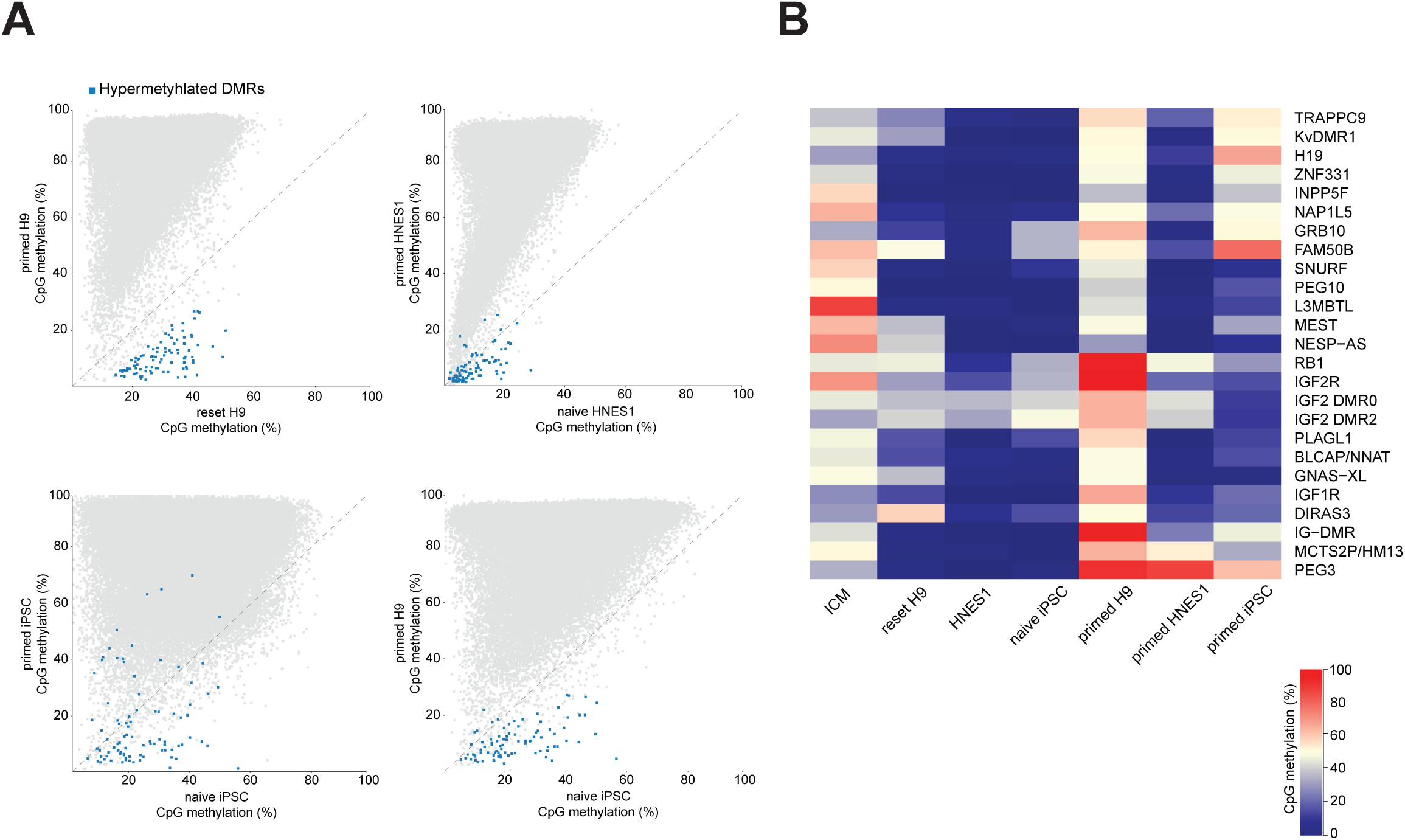
Analysis of CpG methylation globally and at imprinted DMRs. S5A. Scatter plots of CpG methylation percentages over tiles spanning 20 kb. Regions with >10% gain in CpG methylation in reset H9-NK2 cells^9^ compared to conventional primed H9 cells are highlighted in blue in all scatterplots. S5B. Averaged CpG methylation of known DMRs of imprinted maternal and paternal genes.

